# Trial-history biases in evidence accumulation can give rise to apparent lapses

**DOI:** 10.1101/2023.01.18.524599

**Authors:** Diksha Gupta, Brian DePasquale, Charles D. Kopec, Carlos D. Brody

## Abstract

Trial history biases and lapses are two of the most common suboptimalities observed during perceptual decision-making. These suboptimalities are routinely assumed to arise from distinct processes. However, several hints in the literature suggest that they covary in their prevalence and that their proposed neural substrates overlap – what could underlie these links? Here we demonstrate that history biases and apparent lapses can both arise from a common cognitive process that is normative under misbeliefs about non-stationarity in the world. This corresponds to an accumulation-to-bound model with history-dependent updates to the initial state of the accumulator. We test our model’s predictions about the relative prevalence of history biases and lapses, and show that they are robustly borne out in two distinct rat decision-making datasets, including data from a novel reaction time task. Our model improves the ability to precisely predict decision-making dynamics within and across trials, by positing a process through which agents can generate quasi-stochastic choices.

## Introduction

It has long been known that experienced perceptual decision makers deviate from the predictions of optimal decision-theory, displaying several suboptimalities in their decision-making. Among the most pervasive of these is the dependence of behavior on the recent history of observed stimuli, performed actions, or experienced outcomes, despite it being disadvantageous and leading to worse performance (Cho et al. 2002; Gold, Law, et al. 2008; Busse et al. 2011; Carandini and Churchland 2013; Zhang et al. 2014; Fründ et al. 2014; Scott et al. 2015; Abrahamyan et al. 2016; Odoemene et al. 2018; Akrami et al. 2018; Pinto et al. 2018; Urai et al. 2019; Hermoso-Mendizabal et al. 2020; Mendonça et al. 2020; Lak et al. 2020; Mochol et al. 2021; Roy et al. 2021; The International Brain Laboratory et al. 2021; schematized in Fig 1A top). History biases may arise due to a strategy that is optimized for naturalistic settings, where continual learning of priors, action-values, or other decision variables helps agents adapt to changing environments, but is maladaptive in experimental settings where the statistics of the environment are stationary (Yu and Cohen, 2009; Molano-Mazon et al., 2021). To date, decision-theoretic models have accommodated history biases by modeling them as a biasing factor on the perceptual evidence that drives choices (Nevin, 1969; Ratcliff and Rouder, 1998; Bogacz et al., 2006; Busse et al., 2011; Goldfarb et al., 2012; Gardner, 2019; Urai et al., 2019; Hermoso-Mendizabal et al., 2020). In the predominant conceptualization of these models, history biases can be overcome with sufficient perceptual evidence.

**Figure 1:**
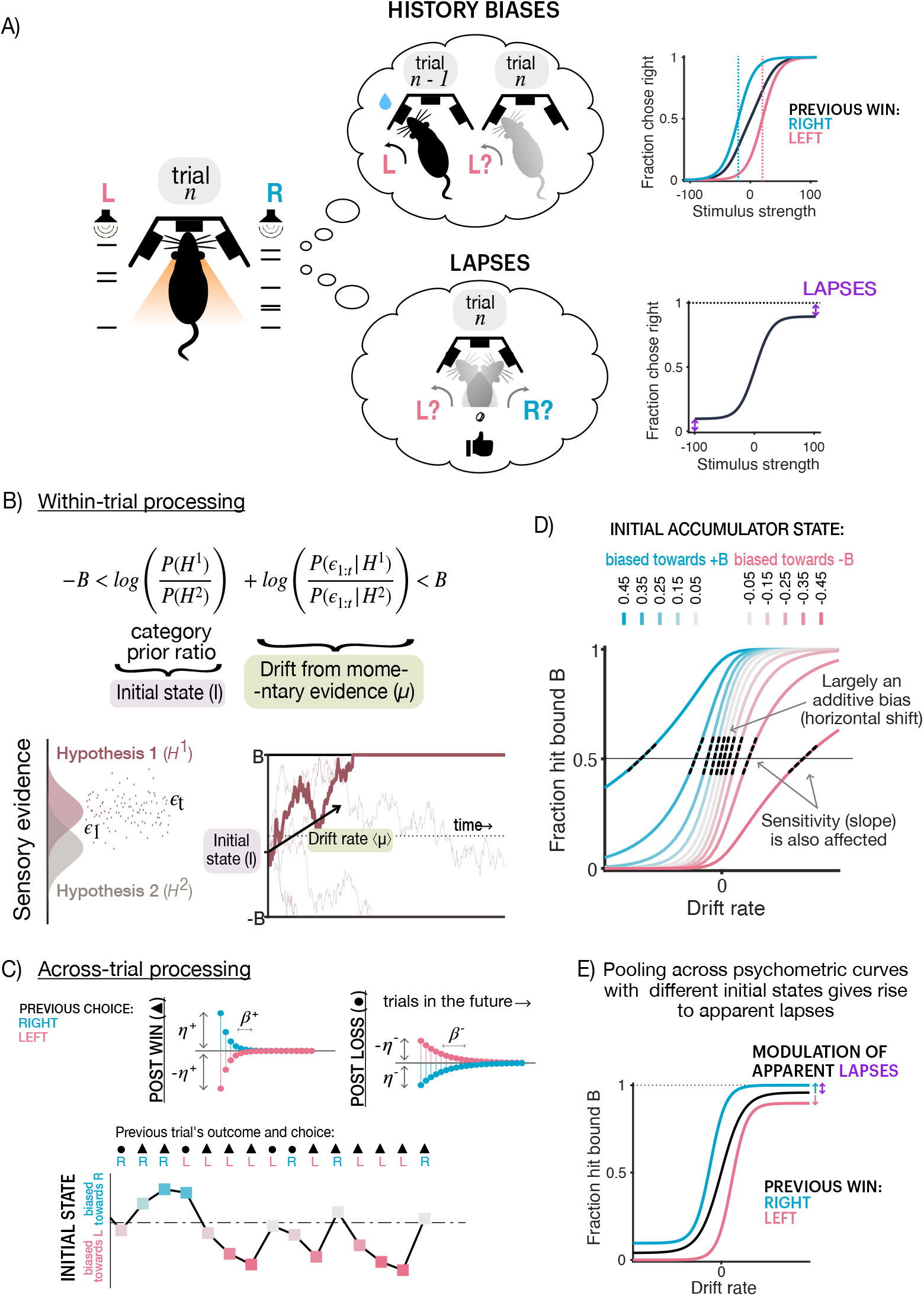
Trial history-dependent initial states give rise to apparent lapses. **(A)** Schematic of two commonly observed suboptimalities in decision-making: history biases (top) and lapses (bottom). (Left): Rat performing a perceptual decision making task, where it has to make one of two decisions (left, right) based on accumulated sensory evidence (auditory clicks on either side). (Top left): History biases i.e. a tendency for the decision on the current trial (n) to be inappropriately influenced by what happened on the previous trial (n-1) in addition to the accumulated sensory evidence. In this example, a previously rewarding leftward decision is likely to be repeated. (Top right): Typically assumed effect of history bias on the psychometric curve, which is the proportion of rightward decisions as a function of the stimulus strength. History biases are thought to most strongly affect decisions when the sensory evidence is weak i.e. around the inflection point of the curve (threshold parameter), shifting it horizontally. (Bottom left): Lapses i.e. a tendency to make seemingly random choices on some trials, irrespective of the accumulated sensory evidence. (Bottom right): Typically assumed effect of lapses on the psychometric curve is vertically scaling the endpoints or asymptotes of the curve. **(B)** Standard normative model of within-trial processing during evidence accumulation. (Top) Decision rule that produces the most accurate decisions in the shortest amount of time, in which a decision is made when the summed log-ratios of category priors and likelihoods exceeds one of two decision bounds. This corresponds to a drift-diffusion process where the prior term sets the initial state (I) and the rate of accumulating evidence sets the drift rate (μ). (Bottom left): Schematic of the generative model, where one of two hypotheses (H1, H2) produce noisy samples of evidence over time (ϵ_t_). (Bottom right): Schematic of the aforementioned drift-diffusion process, showing a sample trajectory based on noisy evidence (bold line) that leads to a rightward decision when the positive bound is hit. Thin lines depict alternate trajectories based on different noisy instantiations of the same drift rate (black arrow). **(C)** Model of across trial processing that can accommodate several forms of prior updates. Past choices and outcomes can additively affect the initial state with different magnitudes (n) and exponentially decaying timescales (*β*) depending on whether they were wins (top left) or losses (top right). (Bottom): Example sequence of trials, labelled by whether they follow a previous win (triangles) or previous loss (circles) on right (R) or left (L) choices, showing the cumulative effect of trial history on initial state updates. Colors denote different initial state biases, same as (C). **(D)** Effect of initial state values on psychometric functions. Colors denote different initial state levels, towards the positive (blue) or negative (pink) decision bounds. Small deviations from 0 in the initial state (grey) lead to largely additive, horizontal biases in the psychometric curve whereas larger deviations (saturated colors) have more complex effects, additionally reducing its effective slope (dotted black lines) or “sensitivity” to the stimulus. **(E)** Pooling different initial state biases gives rise to apparent lapses. Psychometric function (black) pooled across trials with different initial state biases (due to history-based updating) has apparent lapses (purple arrow), moreover conditioning the psychometric curve on whether the previous trial was a rightward (blue) or leftward (pink) win reveals a modulation of these apparent lapses by trial history.

A second widely-recognized but less studied suboptimality is the tendency to “lapse”, or make (asymptotic) errors that are immune to strong evidence (Wichmann and Hill 2001; Law and Gold 2009; Busse et al. 2011; Gold and Ding 2013; Brunton et al. 2013; Carandini and Churchland 2013; Wang et al. 2018; Pinto et al. 2018; Pisupati et al. 2021; Shushruth and Shadlen 2021; schematized in Fig 1A bottom). Because lapses appear to be evidence-independent, they are assumed to arise from nuisance mechanisms that are separate from the perceptual decision-making process and are often imputed to ad-hoc noise sources such as inattention, motor errors etc.

However, several recent results suggest that these two suboptimalities may be linked in their origin. In primates, learning reduces dependence on recent trial history (Gold, Law, et al., 2008) as well as lapse probabilities (Law and Gold, 2009). Intriguingly, mice trained on a visual detection task showed higher levels of history dependence on sessions with higher lapse probabilities (Busse et al., 2011). Moreover, lapses occur in runs (i.e. display Markov dependencies), rather than occurring with the traditionally assumed independent probabilities across trials (Ashwood et al., 2022). Furthermore, lapses have been proposed to reflect forms of exploration (Pisupati et al., 2021) that are sensitive to trial-by-trial updates of variables such as action value. Likewise, neural perturbations of secondary motor cortex and striatum in rodents have been shown to substantially impact both lapses (Erlich, Bialek, et al., 2011; Erlich, Brunton, et al., 2015; Yartsev et al., 2018; Guo et al., 2019; Pisupati et al., 2021; Sindreu et al., 2021) and trial-history influences on decisions (Siniscalchi et al., 2019; Sindreu et al., 2021). Together, these observations challenge the assumption that history biases and lapses have independent causes and raises the possibility that some of the variance ascribed to lapses emerges from history dependence.

We explore the idea that history biases reflect a misbelief about non-stationarity in the world, and demonstrate that normative decision-making under such beliefs gives rise to choices that are both history-dependent and appear to be evidence-independent (i.e. akin to lapses). This corresponds to an accumulation to bound process with a history dependent initial state. We fit this model to a large dataset of choices made by 152 rats trained on an auditory decision-making task. Despite heterogeneity in history biases and lapse rates in this population, we show that the a substantial fraction of lapses can be explained by the presence of history dependence during evidence accumulation. Further, our model predicts the time it takes to make decisions. We test these predictions in a novel task in rats with reaction time reports, and show that it captures patterns of choices, reaction times, and their history dependence. This model significantly improves our ability to predict the temporal dynamics of decision variables within and across trials in perceptual decision making tasks, rendering choices that were previously thought to be stochastic, predictable.

## Results

### A common mechanism produces history biases and apparent lapses

It is often assumed that well trained subjects in two-alternative forced choice (2AFC) tasks have faithfully learnt the likelihood function and priors that determine the structure of the task (Bogacz et al., 2006; Gold and Shadlen, 2007). Under this assumption, the optimal decision making strategy entails combining any knowledge about prior prevalence of available options with the stream of incoming evidence until a desired threshold of confidence is reached in favor of one of the options ^1^ (Gold and Shadlen, 2007; Dayan and Daw, 2008; Drugowitsch, Moreno-Bote, et al., 2012); Fig 1B top). This strategy converges to a drift diffusion model (DDM) when evidence is sampled continuously (Bogacz et al., 2006). In a DDM, one’s belief about the correct option maps onto a diffusing particle that drifts between two boundaries, where the first boundary the particle crosses determines the decision (Fig 1B). Correspondingly, the initial state of this particle encodes the prior belief, and the drift rate is set by the likelihood of incoming evidence (Fig 1B). We refer to the evolving state of the particle in this model as ‘accumulated evidence’.

However, in general, subjects may not know that the task structure is stationary, and might incorrectly assume that it is constantly changing (Yu and Cohen, 2009). In this case, even experienced subjects would not converge to a static estimate of prior probabilities and likelihood functions, but would instead continually update them from trial to trial. Here we consider choice behavior that results from non-stationary beliefs about priors, which result in trial-to-trial updates to the initial accumulator states^2^.

We assume that the initial state of the accumulator (*I*) is set based on the exponentially filtered history of choices and outcomes on past trials. Each unique choice-outcome pair (denoted by *h;* Fig 1C) is tracked by its own exponential filter (i^h^). On each trial *n*, each filter *i^h^* decays by a factor of *β^h^* and is incremented by a factor of *η^h^* depending on the choice-outcome pair on the previous trial:

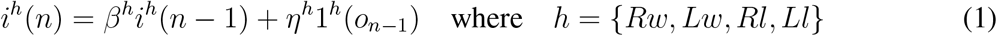

{*Rw, Lw, Rl,Ll*} represent the possible choice-outcome pairs: right-win, left-win, right-loss, and left-loss respectively. *o_n–1_* is the choice-outcome pair observed on trial (n –1) and 1^h^(o_n–1_) is an indicator function that is 1 when *o_n–1_ = h* and is 0 otherwise. The initial state of accumulation, I on trial n is given by the sum of these individual exponential filters:

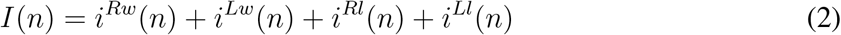

Such a filter can approximate optimal updating strategies under a variety of non-stationary beliefs. As an example, we show that this exponential filter can successfully approximate initial state updates during Bayesian learning of priors under the belief that the prior probabilities of the two hypotheses can undergo unsignaled jumps (Supp Fig. 1; Yu and Cohen 2009; Zhang et al. 2014). Nevertheless, we use this more flexible parameterization to allow for asymmetric learning from different choices and outcomes, which could be beneficial under generative models where one believes that one category persists for longer than another (requiring different decay rates), or correct and incorrect outcomes are not equally informative (requiring different update magnitudes). For instance, in a prior-tracking experiment where previous correct choices had a cumulative effect, but errors had a resetting effect (Hermoso-Mendizabal et al., 2020), this could be captured in the exponential filter by faster decay rates for errors.

What are the consequences of such trial-by-trial updating of initial accumulator states for choice behavior? In a DDM, for a given initial state I and drift rate μ, the probability of choosing the option corresponding to bound *B+* is given by:

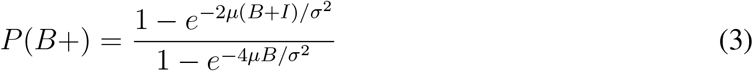

where B is the magnitude of the bound and σ^2^ is the squared diffusion coefficient (derived from Palmer et al. 2005). The resultant psychometric curves for different values of initial accumulator states are plotted in Fig 1D. This expression reduces to a logistic function of *μB/σ^2^* only when I = 0. Small deviations in the initial state largely manifest as additive biases to the evidence, shifting psychometric curves horizontally towards the option favored by the initial state. This corresponds to a change in the psychometric “threshold”, i.e. the x-axis value at its inflection point (Fig 1D lighter colors). Interestingly, large deviations in the initial state produce qualitatively different effects on choices (Fig 1D darker colors). They not only bias the choices towards the option consistent with the initial state but additionally reduce the effective “sensitivity” to evidence. This can be seen as reduction in slope at the inflection point of the psychometric curve (Fig 1D dashed lines). Therefore, trial to trial deviations in the initial state produce history-biased choices which have differently diminished dependence on the evidence.

The average choice behavior obtained by pooling choices with different history-biased initial states is a mixture of psychometric curves with varying thresholds and sensitivity to perceptual evidence. Such a psychometric curve is heavy-tailed (Shen and Ma, 2019; Nguyen, Josić, et al., 2019) and appears to have asymptotic errors or “lapse rates” (Fig 1E, black curve). These asy mptotic errors are not truly evidence-independent, random decisions or *true lapses*, rather they are *“apparent lapses”* arising from evidence accumulation with deterministic history-based updates to the initial accumulator state. In such a setting, the psychometric curves obtained by conditioning on past trials’ choice and outcome, or *history conditioned* psychometric curves, are both horizontally and vertically shifted, i.e. they show history-dependent modulations in both threshold and lapse rate parameters (Fig 1E, Supp Fig. 2B). In this formulation, trial-history modulated lapse rates are uniquely produced by history-biased initial accumulator states (and therefore reflect apparent lapses), in contrast to lapse rates observed in the unconditioned psychometric curve which might have additional extraneous causes (Wichmann and Hill, 2001; Ashwood et al., 2022; Pisupati et al., 2021), and therefore reflect both apparent and true lapses.

In this model, because history modulations of psychometric thresholds and lapse rates arise from one unified process, they are not allowed to vary independently of the decision-making process, or of each other. Rather their relative magnitudes are intimately coupled with and constrained by accumulation variables. For instance, increased magnitudes or timescales of initial state updating produce large fluctuations in the initial accumulator state across trials. This in turn reduces the effective sensitivity of the accumulation process to evidence, giving rise to more apparent lapses and history biases (Supp Fig. 2A). Similarly, changes in within-trial parameters of accumulation can dramatically influence these history modulations (Supp Fig. 2C). Decisions made with smaller accumulator bounds are more sensitive to initial state modulations, and therefore give rise to more apparent lapses and higher modulations of lapse rates and thresholds. Higher levels of sensory noise have a similar effect, yielding more apparent lapses, consistent with recent reports of lapse rates being modulated by sensory uncertainty (Pisupati et al., 2021). Finally, impulsive integration strategies that overweigh early evidence rather than accumulating uniformly (Bogacz et al., 2006) exaggerate the influence of initial states, producing more apparent lapses and history biases.

Some definitions:

- *Lapse rate:* Lapse rates capture the difference between perfect performance and observed performance at the asymptotes, measured through sigmoidal fits to the psychometric curves.
- *True lapse:* A true lapse is an independent cognitive process through which agents generate stochastic evidence-independent choices.
- *Apparent lapses:* Apparent lapses are deterministic evidence-dependent choices, that nonetheless contribute to lapse rates.

### Rats display varying degrees of history-dependent threshold and lapse rate modulation

We sought to test if the co-modulations posited by our model are present in rat decision-making datasets, in order to ascertain whether a unified explanation could underlie the links between history biases and lapses.

We first examined whether and how rat decision-making strategies were affected by trial history. We analyzed choice data from 152 rats (37522 ±22090 trials per rat, mean ±SD; Supp Fig.3A) trained on a previously developed task that requires accumulation of pulsatile auditory evidence over time (‘Poisson Clicks’ task, Brunton et al. 2013). In this task, the subject is presented with two simultaneous streams of randomly-timed discrete pulses of evidence, one from a speaker to their left and the other to their right (Fig 2A). The subject must maintain fixation throughout the stimulus, and subsequently orient towards the side which played the greater number of clicks to receive a water reward. The trial difficulty, stimulus duration, and correct answer were set independently on each trial. Because this task delivers sensory evidence through randomly but precisely timed pulses, it provides high statistical power to characterize decision variables that give rise to the choice behavior.

**Figure 2:**
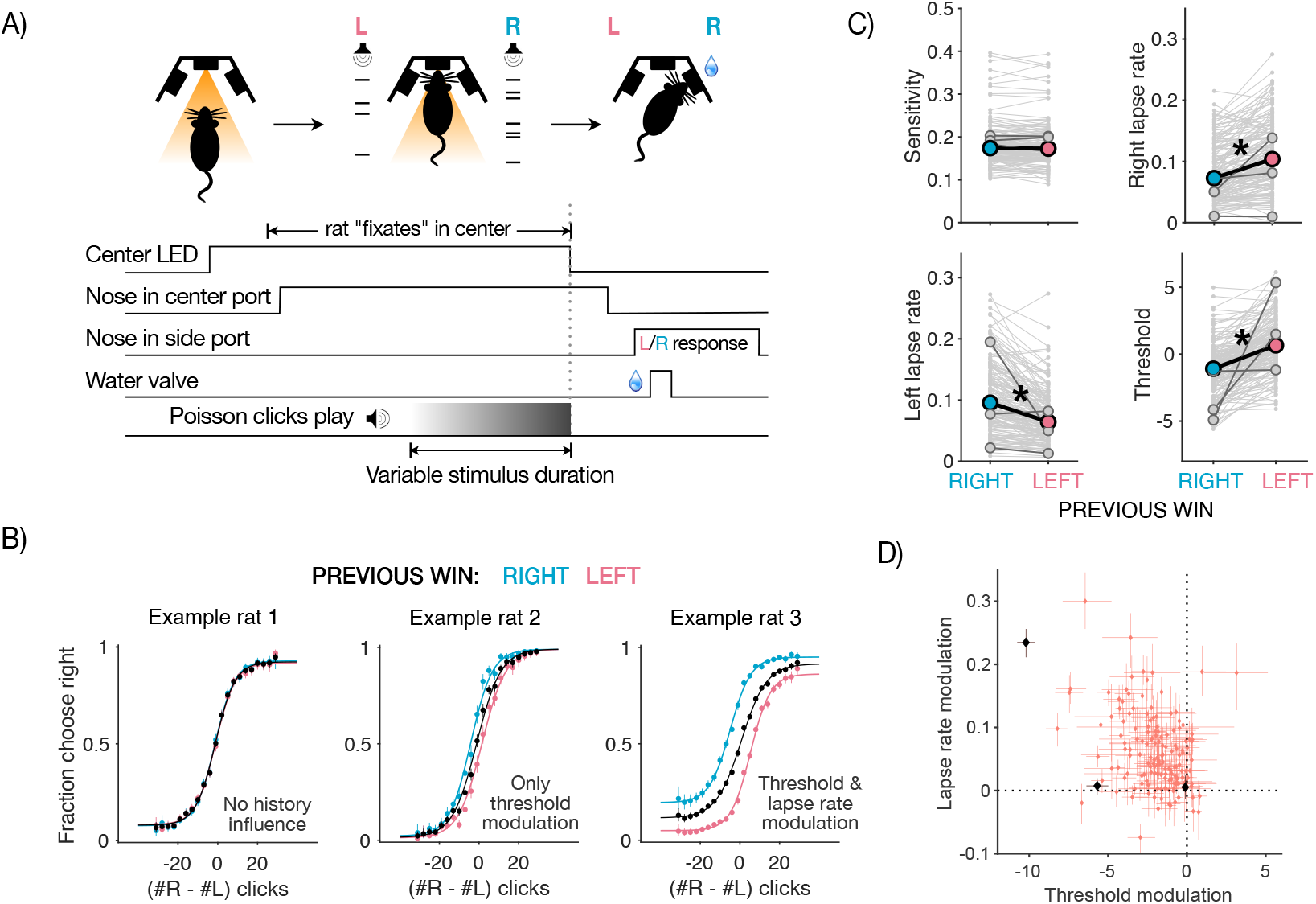
History-dependent threshold and lapse rate modulations in a large-scale rat dataset. **(A)** Schematic of evidence accumulation task in rats: (Top): Phases of the ‘Poisson clicks’ task, including trial initiation in center port (left), evidence accumulation based on two streams of Poisson-distributed auditory clicks (middle) and choice report in one of two side ports followed by water reward for correct choices (right). (Bottom): Time-course of trial events in a typical trial. **(B)** Individual differences in history-dependence: Psychometric functions of three example rats from a large-scale dataset, displaying different kinds of history modulation. Choices are plotted conditioned on previous left (blue), right (pink) or all wins (black). (Left): Example rat with no history-dependence in choices, resembling the ideal observer. (Middle): Example rat with modulations of the threshold parameter alone, resembling the dominant conceptualization of history bias. (Right): Example rat with history-dependent modulation of both threshold and lapse rate parameter, similar to the majority of the population. **(C)** Dataset displays significant modulations of both threshold and lapse rate parameters: Scatters showing parameters of psychometric functions following leftward wins (post left, blue) or rightward wins (post right, pink). Each pair of connected gray points represents an individual animal, solid colored dots represent average parameter values across animals. Trial history does not significantly affect the sensitivity parameter (top left) but significantly affects left, right lapse rate and threshold parameters (top right and bottom panels). **(D)** Scatter comparing threshold and lapse rate modulations in the entire population. Each dot is an individual animal, error bars are ±95% bootstrap CIs. Black points represent example rats. The majority of the population lies in the top left quadrant, showing co-modulations of both threshold and lapse rate parameters by history.

Rats performed this task accurately (0.79 ± 0.04, mean accuracy ± SD, Supp Fig.3B). Performance was stable with little to no change in accuracy across trials (mean slope ± SD across rats of linear fit to hit rate over trials: 1.13×I0^-7^± 8.90×I0^-7^; Supp Fig.3C) reflecting asymptotic behavior rather than task acquisition. Rats showed history dependence in their choices, largely tending towards a “win-stay, lose-switch” dependence (Supp Fig.3E). We found substantial individual variability in the dependence of rats’ choices on history in the dataset. Some rats were weakly influenced by history (Fig 2B left) while others showed a history-dependent modulation of the psychometric threshold parameter (Fig 2B middle) or a history-dependent modulation of both threshold and lapse rate parameters (Fig 2B right). The population as a whole most closely resembles the Example rat 3, with both threshold and lapse rate parameters being significantly different following left and right wins while sensitivity is not affected (*p =* 0.8 for sensitivity, 3 × 10^-17^ for bias, 8 × 10^-8^ for left lapse, 6 × 10^-7^ for right lapse, Mann-Whitney U-test, Fig 2C). As predicted by our model (Fig 1E), trial-history biased both threshold and lapse rate parameters in the same direction (e.g. both biased toward rightward choices following right rewards). Moreover, the vast majority of rats show co-modulations of both parameters by history (Pearson’s correlation coefficient: r = −0.35, p = 7.28 × 10^-6^; Fig 2D). Across rats, on average 17 ±12%of total lapse rates are modulated by trial history and therefore could potentially reflect apparent rather than true lapses (Supp Fig.3D). These findings support the conclusion that rat decision-making strategies, while idiosyncratic, largely show history-dependent effects consistent with our model. Next, we tested the model more directly using trial-by-trial model fitting.

### History-dependent initial states capture comodulations in thresholds and lapse rates in the data

To test whether the observed history modulations in thresholds and lapse rates arise from trial-by-trial updates to the initial accumulator state, we extended an accumulator model previously adapted to this pulsatile task (Brunton et al., 2013) to incorporate History-dependent Initial States (abbreviated as HISt, Fig 3A). As before, we model this history-dependence using an exponential filter over past trials’ choices and outcomes (Fig 1C). Hence, across trials the accumulator model with HISt produces apparent lapses, as well as coupled history modulations in psychometric threshold and lapse rate parameters.

**Figure 3:**
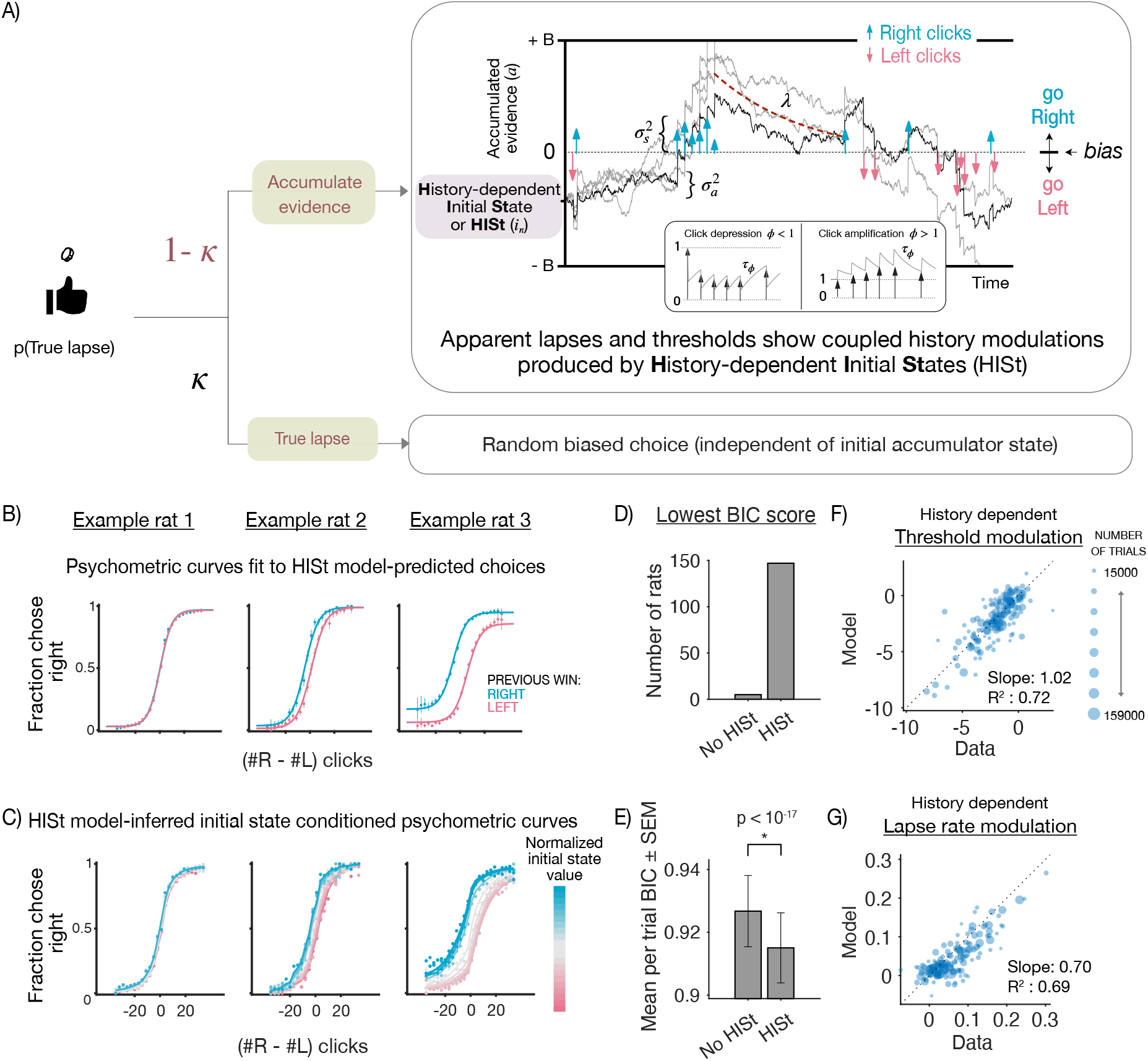
>History-dependent initial states capture comodulations in thresholds and lapse rates in the data. **(A)** Schematic of the model (accumulator with HISt) used to fit rat data in the Poisson Clicks task. **(Top)**: Schematic of accumulation-to-bound model whose initial states are modulated by trial history producing history-dependent apparent lapses and threshold modulations. The model consists of sensory noise 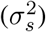 in click magnitudes, adaptation of successive click magnitudes based on an adaptation scale (*φ*) and timescale (*Tψ*), accumulator noise 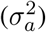 added at each timestep, leak in the accumulator (λ), and decision bounds +/-B. We refer to this accumulator model with History-dependent Initial States as ‘HISt’ **(Bottom)**: On κ fraction of trials, the model occasionally chooses a random action irrespective of the initial state and stimulus, with some bias (ρ) reflecting a motor errors or random exploration. These true lapses are not modulated by history, such that any history modulations arise from the initial states alone. **(B)** Model fits to individual rats: Psychometric data from 3 example rats conditioned on previous rightward (blue) or leftward (wins), overlaid on model-predicted psychometric curves (solid line) from the accumulation with HISt model. The three example rats were chosen to illustrate the diversity of history effects in the dataset, ranging from no history effects (left) - to history effects that largely created horizontal biases (center) and history effects that additionally affected lapse rates (right). **(C)**: Psychometric curves (solid line) from the same example rats conditioned on model-inferred initial states (colors from pink to blue), showing a similar pattern to analytical predictions in Fig 1D. **(D)** Distribution of best fitting models for individual rats: Overall bar height for each model denotes the total number of rats for which that model scored the lowest BIC score. **(E)** Model comparison using BIC by pooling per trial BIC score across rats and computing mean. Lower scores indicate better fits. Mean of per trial BIC scores across rats were significantly lower for model with HISt (p = 9.85 × 10^-18^, paired t-test). Error bars are SEM. **(F)** Individual variations in history modulations captured by the accumulator model with HISt: History modulations of threshold parameters measured from psychometric fits to the raw data (x-axis) v.s. model predictions (y-axis). Individual points represent individual rats, point sizes indicate number of trials. The model captures a majority of the variability as evidenced by the points lying close to the unity line. **(G)** same as (F) but for history dependent lapse rate modulations. The model captures a majority of the variability in lapse rate modulations, implying that the magnitude of threshold and lapse rate modulations are coupled as predicted by our model, and that history-dependent initial accumulator states contribute to apparent lapses in this dataset.

Within a trial, our accumulator model leverages knowledge of the timing of each evidence pulse to model the sensory adaptation process as well as to estimate the noise and drift of the accumulator variable (Fig 3A top bubble, Methods). The model includes a feedback parameter that controls whether integration is leaky, perfect, or impulsive. Following Brunton et al. 2013,this model also includes (biased) random choices independent of the accumulator value on a small fraction of trials (κ) - we consider decisions arising from this process to be “true lapses” because they are evidence independent, unlike apparent lapses which still retain some evidence-dependence (Fig 3A bottom bubble).

We performed trial-by-trial fitting of the accumulator model with and without History-dependent Initial States (HISt) to choices from each rat using maximum likelihood estimation (Methods). We find that the accumulator model with HISt captures both psychometric curve threshold and lapse rate modulations well across different regimes of rat behavior, as evident from fits to example rats (Fig 3B). Moreover, conditioning rats’ psychometric curves on model-inferred initial state values reveals that the initial state captures a large amount of variance in choice probabilities (Fig 3C), resembling theoretical predictions (Fig 1C). This shows that the initial state is a key explanatory variable underlying choice variability both across and within individuals, that jointly modulates multiple features of the empirical psychometric curves in a parametric fashion. We used Bayes Information Criterion (BIC) to determine whether adding HISt to the accumulator model was warranted (Fig 3D-E). Individual BIC scores recommended that adding HISt was warranted in 147/152 rats (Fig 3D). This model also best captured choices across the population as a whole, with significantly lower mean BIC scores across rats (Mean per trial BIC score for HISt: 0.91 ±0.01 vs no HISt: 0.93 ±0.01, p = 9.85 × 10^-18^, paired t-test; Fig 3E). Next, we compared the psychometric threshold and lapse rate modulations produced by this model to the modulations in the data, as determined by conditioning the psychometric functions on trial-history (Fig 3B). As predicted, the model successfully accounted for modulations in both these distinct psychometric features via the singular process of trial-by-trial history-dependent updates to the initial accumulator state. Across individuals, the model with HISt captured a substantial amount of variance [R^2^ = 0.72 (threshold parameter), R^2^ = 0.69 (lapse rate parameter)] and showed good correspondence to the empirical modulations in data [slope= 1.02 (threshold parameter), slope = 0.70 (lapse rate parameter)].

In our model, apparent lapses show history modulations since they are produced by history-dependent initial accumulator states, while true lapses do not since they result from an occasional flip in the final choice and are independent of the accumulator value (following Brunton et al., 2013). Such kinds of true lapses could reflect errors in motor execution or random exploratory choices made despite successful accumulation (Supp Fig 4B). However true lapses could also occur due to inattention, i.e. an occasional failure to attend to the stimulus. In such cases, the optimal strategy devoid of sensory evidence is to deterministically choose the side favored by the initial accumulator state (Supp Fig 4C). Therefore, inattentional true lapses, while remaining evidence independent, may nevertheless be modulated by history due to their initial state dependence. In order to account for this possibility, we fit an additional “inattentional” variant of the accumulator model with HISt (Supp Fig 4A,C), and found that it was closely matched on BIC scores with the previous model which we label as the “motor error” variant (Supp Fig 4E,F). Moreover, the inattentional variant, which additionally allows true lapses to depend on history, only captured slightly more variance in history modulations of lapse rates, at the expense of history modulations of thresholds (Supp Fig 4D). These minor improvements support the hypothesis that apparent lapses produced by history-dependent initial states (rather than true lapses due to motor error or inattention) are the major driver of history-dependent co-modulations in psychometric thresholds and lapse rates in the dataset.

To summarize, our model predicted that the initial accumulator state should be the underlying variable that jointly drives history-dependence in thresholds and lapse rates – implying that our accumulator model with HISt should be able to simultaneously capture variability in both these parameters across rats. Our rat dataset strongly supports this prediction, lending evidence to the hypothesis that history-dependent initial states give rise to apparent lapses, and are the common cognitive process that underlie links between these two suboptimalities that were previously thought to be distinct from each other.

### Reaction times support history-dependent initial state updating

In our model with history-dependent initial accumulator states, the time it takes for the accumulation variable to hit the bound determines the duration that the subject deliberates for, before committing to a choice. Therefore in addition to choices the model makes clear predictions about subjects’ reaction times (RTs). We sought to test if these predictions are borne out in subject RTs.

To this end, we trained rats (n = 6) on a new variant of the auditory evidence accumulation task (Fig 2A), with two key modifications that allowed us to collect reaction time reports (Fig 4A). First, in this new task the stimulus is played as long as the rat maintains their nose in the center port (or “fixates”) and stops immediately when this fixation is broken. Second, in this task the rat has to correctly report which speaker’s auditory click train is sampled from a higher Poisson rate to receive a water reward (unlike the non-reaction time task where subject has to report the side which played the greater number of clicks). Rats perform this task with high accuracy (Fig 4B left panel, average accuracy: 0.75 ±0.02, number of trials 37205 ±14247, mean ±SD). Similar to the previously analyzed data, their choices are impacted by recent trial history (Fig 4B right panel). Moreover, trial-history dependent modulation of psychometric function parameters (Fig 4C) resembles that of the non-reaction time task (Fig 2C; *p* = 0.69 for sensitivity, 0.004 for threshold, 0.02 for left lapse rate, 0.02 for right lapse rate, Mann-Whitney U-test). Once again, this history modulation of both psychometric threshold and lapse rate parameters in tandem is consistent with our singular accumulator model with history-dependent initial states.

**Figure 4:**
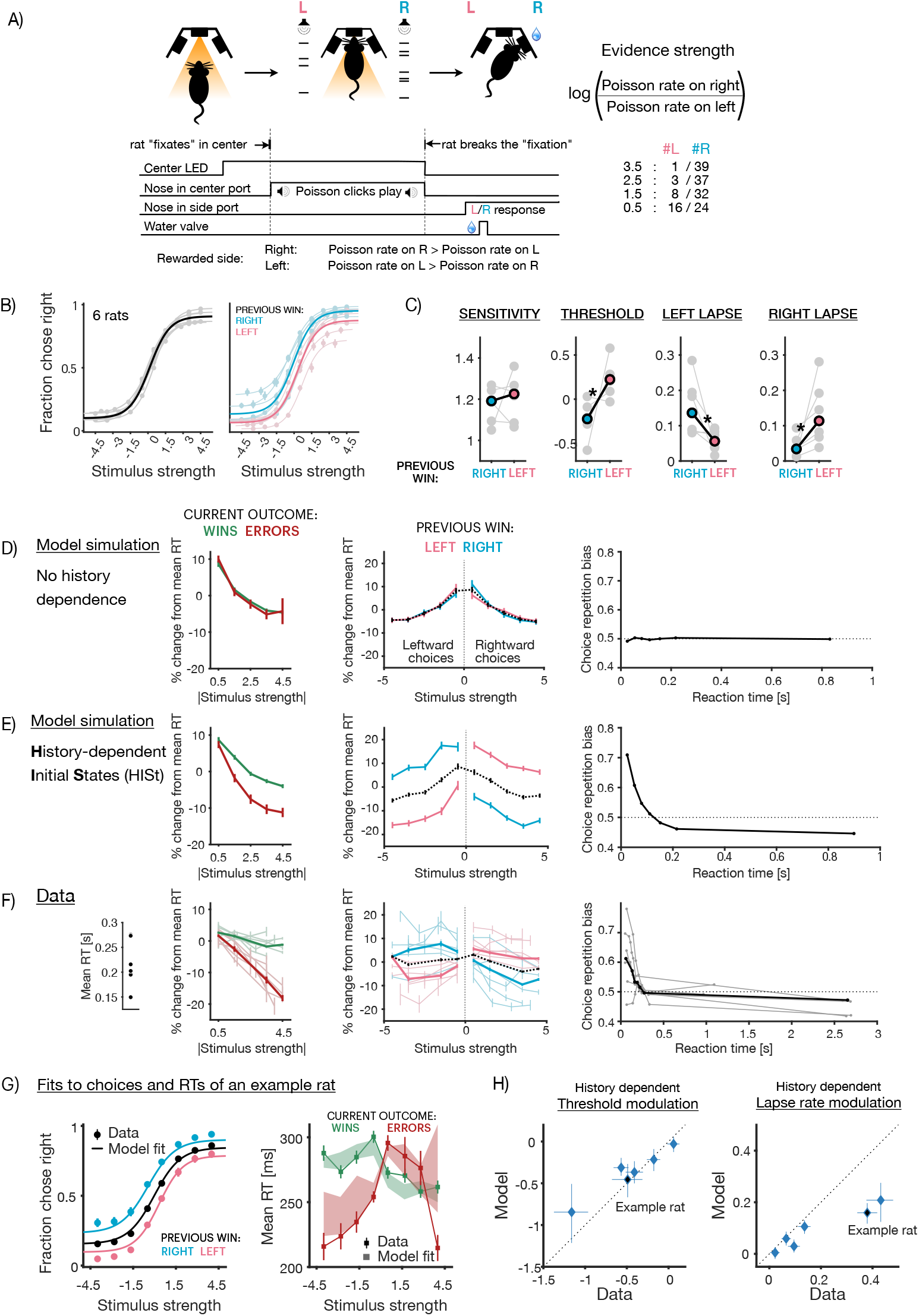
Model predictions about reaction times are borne out in data. **(A)** Schematic of reaction time task in rats, with similar structure to (Fig 2A), with two modifications: rats are allowed to break “fixation” anytime during the trial and make a choice, and are rewarded for choosing the side with the higher Poisson rate, encouraging longer sampling for more accurate estimates. **(B)** Average choice behavior on all trials (left) and following previous right or left wins (right) of 6 rats on the reaction time task (solid line), overlaid on individual rat behavior (translucent lines) **(C)** Average parameters (solid points) of history-conditioned psychometric curves, overlaid on individual parameters (translucent points) showing significant history modulations in threshold and lapse rate parameters (*p* 0.05, Mann-Whitney U-test) **(D-F)** Reaction time signatures D) expected from accumulator models with no history dependence in initial states, E) expected from accumulator models with history dependent initial states and F) observed in data. First, error reaction times are expected to be shorter if initial states are history dependent, as seen in data (Left column, red curves are below green curves in E,F). Second, reaction times on trials following right wins are expected to be lower on rightward stimuli (positive half of x-axis), and similarly following left wins (Middle column, blue (pink) curves on the right (left) are lower than dotted lines in E,F). Finally, repetition biases in choices are expected to occur more frequently for short reaction times, when the effect of initial states is strong (Right column, curves are above dotted line for smaller RTs in E,F). **(G)** Joint fits of the accumulator model with history-dependent initial states to choices (left) and reaction times (right) of an example rat show good correspondence to data. Data represented by points (circles: choices, squares: reaction times) and model fits represented by lines (choices) or shaded bars (reaction times, thickness represents 95% bootstrap prediction intervals). Reaction times (right) are split by wins (green) or errors (red). **(H)** Scatter plot showing correspondence between history modulations in threshold (left) or lapse rate (right) parameters derived from data (x-axis) and model fits (y-axis). Individual points represent individual rats.

Moreover, reaction times (RTs) of these rats display several signatures predicted by our model (Fig 4D-F). First, trial-to-trial variability in the initial state of the accumulator is expected to give rise to shorter RTs on error trials compared to correct trials (Fig 4E, left; Ratcliff and Rouder 1998). This is because trials in which the initial state is closer to the incorrect bound are more likely to be errors, but because of the closer bound they are also likely to hit it faster. This is unlike a standard DDM with no trial-to-trial variability in parameters, where RTs for correct and error trials are of similar magnitudes (Fig 4D, left). Indeed in the rat dataset, error RTs are consistently shorter than correct RTs across rats (Fig 4F, left). Second, initial state updates towards previously rewarded choices (such as in a win-stay agent) are expected to produce shorter RTs when the current stimulus favors the previously rewarded choice (Fig 4E, middle; Yu and Cohen 2009; Goldfarb et al. 2012). We find that this signature is also present in the dataset across rats (Fig 4F, middle). Finally, variability in the initial state is most influential early in the decision process, predicting that the majority of history dependence in choices occurs on trials with fast RTs (Fig 4E, right; Urai et al. 2019). Indeed, the data displays this pattern as well, with repetition bias being most prominent for short RTs, disappearing and turning into a weak alternation bias for long RTs (Fig 4F, right). Taken together, these three signatures offer strong, complementary evidence from RTs for the prevalence of history-dependent initial states in rats performing this evidence accumulation task.

We directly test if our model can simultaneously capture reaction time patterns and history-modulation of psychometric threshold and lapse parameters by jointly fitting choices and RTs of individual subjects in a trial-by-trial fashion (see Methods). We find that the history-dependent initial state model jointly captures patterns of choices, reaction times, and their history modulations in the data (Fig 4G - fits from example rat, Supp Fig. 5 - fits from all rats). This model accounts for substantial variance in history-dependent threshold and lapse rate modulations (Fig 4H). We also fit a hybrid variant of the accumulator model with HISt that flexibly allows true lapses to be motor-error like and unaffected by history, or inattention-like and additionally be modulated by history (Supp Fig. 6A,B). While this model has a better BIC and leads to a slight improvement in correspondence to the history modulation of psychometric lapse rates, it does so at the cost of correspondence to modulations in psychometric thresholds (Supp Fig. 6C-E), once again largely implicating HISt and its resultant apparent lapses (rather than true lapses) in the co-modulation of both parameters.

Overall, these results show that the history-dependent initial state updates that we invoked to explain apparent lapses in rodent data are corroborated by their reaction times, and accounting for them can help render a sizable fraction of decisions — that would have been otherwise attributed to noise — more predictable both within and across trials.

## Discussion

History biases and lapses have both long been known to impact perceptual decision-making across species. However, they have largely been assumed to be distinct from each other, despite their frequent co-occurrence and co-modulation. Here, we propose that normative accumulation under misbeliefs of non-stationarity can produce both history biases and apparent lapses, offering an explanatory link between the two suboptimalities. This corresponds to history-dependent trial-to-trial updates to the initial state of an evidence accumulator. We show that such updates produce choices with varying biases in psychometric thresholds as well as varying sensitivities to evidence, yielding apparent, history-modulated lapse rates when choices are averaged across trials (Fig 1). Our model postulates that the initial state of the accumulator is a key underlying variable that jointly modulates psychometric thresholds and lapse rate parameters, with the exact nature of this comodulation determined by the within and across trial parameters governing evidence accumulation. We tested this model in a large rat dataset consisting of choices from 152 rats (Fig 2) and confirmed its predictions using detailed model-fitting. We found that the singular process of history-dependent initial states successfully captured a substantial amount of variance in history modulations of both thresholds and lapse rates in the dataset (Fig 3). Finally, we tested the reaction time predictions of the model in a novel task in rats, and confirmed that the data showed signatures of initial state updating. The model could successfully capture choices, reaction times, and history modulations in psychometric thresholds and lapse rates (Fig 4). Altogether, our results suggest that history biases and a substantial amount of variance attributed to lapses may reflect a common mechanistic process, whose evolution can be precisely tracked both within and across trials.

History biases in perceptual decision making tasks have been modeled using initial state updates to DDMs in humans and non-human primates (Gold, Law, et al., 2008; Goldfarb et al., 2012; Zhang et al., 2014). These studies tended to have relatively small magnitudes of history bias, and miniscule lapse rates, hence being well captured by small deviations in the initial state of a DDM, which largely yield horizontal shifts in the psychometric function. This regime of initial state updates is well approximated by a logistic function with additive biases, which is the dominant descriptive model used to characterize history-dependent psychometric curves (Busse et al., 2011; Carandini and Churchland, 2013; Fründ et al., 2014; Abrahamyan et al., 2016; Gardner, 2019; Pinto et al., 2018; Odoemene et al., 2018; Urai et al., 2019; Hermoso-Mendizabal et al., 2020; Roy et al., 2021; Ashwood et al., 2022; Bolkan et al., 2022). However, as we demonstrate, when deviations in the initial state are large, this logistic approximation breaks down. This fact has been overlooked in much of the literature. Consequently, even in datasets with large history biases and lapses, the logistic formulation continues to be favored (Odoemene et al., 2018; The International Brain Laboratory et al., 2021; Roy et al., 2021; Ashwood et al., 2022), albeit requiring additional components. Our demonstration predicts that the full range of initial state effects should resemble concurrent, trial-by-trial changes in both threshold and sensitivity parameters of the logistic function. Indeed, Ashwood et al. 2022 found that apparent lapses in several rodent datasets can be better captured by runs of trials with such concurrent modulations, yielding biased “disengaged” states.

In our treatment, we only considered history-dependent updates to the initial state of a DDM. Such a mechanism is normative under non-stationary beliefs about the prior ^3^, which is our favored interpretation as it aligns with other studies of history biases (Gold, Law, et al., 2008; Yu and Cohen, 2009; Summerfield and Koechlin, 2010; Goldfarb et al., 2012; Mulder et al., 2012; Abrahamyan et al., 2016; Molano-Mazon et al., 2021). Nevertheless, these updates may also reflect other heuristic strategies (Gigerenzer and Gaissmaier, 2011) which we accommodate using our flexible parameterization of initial state updates. Animals may entertain non-stationary beliefs about other elements of the decision process, such as the rewards or likelihoods (Dayan and Daw, 2008; Mendonça et al., 2020; Lak et al., 2020; Pisupati et al., 2021). Normative updating in such situations still reduces to initial state updates in simple settings (for e.g. non-stationary rewards for a single difficulty; Simen et al. 2009; Rorie et al. 2010), but in more complex ones it *additionally* affects drift rates (Palmer et al., 2005; Eckhoff et al., 2008; Hanks, Mazurek, et al., 2011; Drugowitsch, Mendonça, et al., 2019; Fan et al., 2018; Urai et al., 2019; Mendonça et al., 2020). Trial-to-trial variability in drift rates is known to give rise to longer error RTs than correct RTs (Ditterich, 2006a; Ditterich, 2006b; Drugowitsch, Moreno-Bote, et al., 2012; Nguyen and Reinagel, 2020), which is a signature often reported in monkeys and humans (Roitman and Shadlen, 2002; Shevinsky and Reinagel, 2019). Although we don’t see this reaction time signature of drift rate variability in our dataset – instead we see signatures of initial state variability, with error RTs being shorter than correct RTs, rather than longer – drift rate updates may be another potential mechanism by which history-modulated apparent lapses could arise.

Lapse rates are often considered to be a mixed bag comprising several different noise processes, yet most studies so far have focused on one or more of these component processes in isolation (Pisupati et al., 2021; Ashwood et al., 2022). In this work, we have attempted a more expansive approach of considering multiple processes at once, in an attempt to partition lapse rate variance into mixtures of deterministic and stochastic components. We distinguished apparent lapses that interact with sensory evidence from two models of “true” lapses that are both evidence independent — motor error or exploration, which does not interact with the accumulator, and inattention, which may still depend on its initial state. While we find that the behavior of our rats is best described by a mixture of apparent lapses and the latter two true lapse variants, it is primarily the apparent lapses (rather than either true lapse variant) that captures the links between the sub-optimalities i.e. the history-dependent comodulations in psychometric thresholds and lapse rates.

A previous study proposed an evidence-dependent model of true lapses, uncertainty-guided exploration (Pisupati et al., 2021), in order to account for the scaling of lapse rates with sensory noise. Although we don’t explicitly consider this model, our model of apparent lapses already displays this property, with higher levels of sensory noise leading to more frequent apparent lapses.

Our model predicts that an increased reliance on history (i.e., larger shifts of the initial states) should produce more apparent lapses. Indeed, this could provide an explanation that links disparate sets of observations from previous studies: while some studies have reported that perturbations of secondary motor cortex and striatum give rise to higher lapse rates (Erlich, Brunton, et al., 2015; Yartsev et al., 2018; Guo et al., 2019; Sindreu et al., 2021; Pisupati et al., 2021), others have shown that the effects of perturbing these regions seems to resemble an increased history-dependence (Sindreu et al., 2021; Luo et al., 2021). Interpreting these results through the lens of our model, we’d conclude that these regions play a crucial role in the interaction of history-dependent initial states with sensory evidence. Indeed, Luo et al. 2021 find that this increased history dependence upon perturbation is mediated by increased bias in initial value of the neurally derived accumulator variable. Our model could also help explain why Busse et al.2011 found that mice with higher lapse probabilities showed higher history dependence, or results from The International Brain Laboratory et al. 2021 who observed a modulation in lapse rates in addition to horizontal biases upon explicit manipulation of category priors.

One interesting future line of investigation is to probe the precise nature of the model of non-stationarity over priors assumed by animals in such tasks. The range of parameter values inferred using our flexible formulation could offer a useful starting point for this line of investigation. For instance, Dynamic Belief Models (Yu and Cohen, 2009; Ryali et al., 2018), a popular class of generative models over priors, correspond to a narrowly constrained set of parameter values in our model. Such an understanding would not only afford more reliable control of behavior and more accurate interpretation of neural correlates in stationary tasks, but could also yield insight into the inductive biases that allow animals to learn quickly and efficiently in non-stationary, naturalistic settings.

## Methods

### Subjects

Animal use procedures were approved by the Princeton University Institutional Animal Care and Use Committee (IACUC #1853). All subjects were adult male Long Evans rats, typically housed in pairs. Rats that trained during the day were housed in a reverse light cycle room. Rats had free access to food but in order to to motivate them to work for water reward, they were placed on a controlled water schedule: 2-4 hours per day during task training, usually 7 days a week and between 0 and 1 hour ad lib following training.

### Drift diffusion model of decision-making

We use a standard formulation of sequential decisionmaking based on (Bogacz et al., 2006; Drugowitsch, Moreno-Bote, et al., 2012), in which an agent is faced with a stream of noisy sensory evidence *ϵ*_1:*t*_ coming from one of two hypotheses H_1_ and H_2_. The agent has to decide between sampling for longer or choosing one of two actions *L,R* (reaction time regime) or has to choose one of two actions after a fixed amount of evidence (fixed duration regime). Such a problem can be formulated as one of finding an optimal policy *π_t_* in a partially-observable markov decision process (Rao, 2010; Drugowitsch, Moreno-Bote, et al., 2012), whose solution can be written as a pair of thresholds on the log-posterior ratio 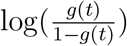, where g(*t*) = *p*(*H*_1_|*ϵ*_1:*t*_:

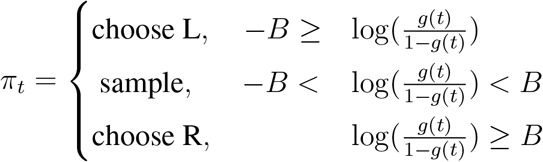

The log posterior ratio can be further broken down into a sum of log prior ratios and log likelihood ratios, using Bayes rule:

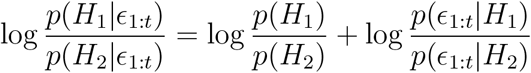

The optimal policy can equivalently be expressed in terms of the prior and sum of momentary sensory evidence *x(t) = ∑_t_ ϵ_t_*, which are sufficient statistics of the posterior (Drugowitsch, Moreno-Bote, et al., 2012; Piet et al., 2018). In the continuous time limit, when the average rate of evidence increments or drift rate is μ, and the standard deviation of sensory noise is σ, this corresponds to a drift diffusion model that terminates when it reaches one of two bounds (Bogacz et al., 2006) and whose initial state *I* is proportional to the log prior ratio:

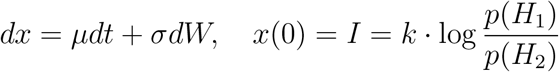

In this case, the probability of choosing rightward actions, i.e. hitting the upper bound can be written analytically as follows (derived from Palmer et al. 2005):

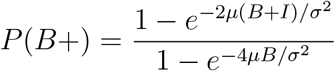

In cases where trial difficulties (and hence drift rates) vary from trial to trial the optimal policy includes time-dependent, collapsing bounds on the posterior. However, under certain circumstances, constant bounds on *X_t_* ∑_*t*_ ϵ_*t*_ implement close-to-optimal collapsing bounds on the posterior (Denève, 2012; Drugowitsch, Moreno-Bote, et al., 2012), which is the regime we assume for our analysis.

### Models of initial state updating

We model initial state updating as a sum of exponential filters over past choice-outcome pairs (*Rw:* right-wins, *Lw:* left-wins, *Rl:* right-loss, *Ll:* left-loss). So the initial state *I* at trial *n* + 1 is given by:

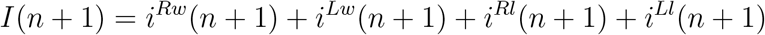

where each filter *i^h^* decays by a factor of *β^h^*, and is incremented by a factor of *η^h^* following the observation of that particular choice-outcome pair, i.e

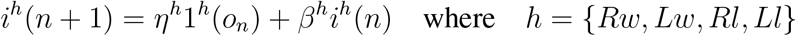

*o_n_* is the choice-outcome pair observed on trial n and 1^h^(o_n_) is an indicator function that is 1 when *o_n_* = *h* and is 0 otherwise.

When *β^h^* and *η^h^* are the same *∀h*, this rule reduces to an approximation of the Bayesian update for the Dynamic Belief Model (Yu and Cohen, 2009), which tracks a prior that undergoes discrete unsignaled switches at a fixed rate. We compared this unconstrained model to models with various constraints on the decay and magnitude parameters (same parameters for corrects v.s. errors, left v.s. right etc). While model comparison revealed that not every rat required all parameters to be different, the unconstrained model is the most general form that best captures behavior across rats.

### Psychometric curves

Psychometric curves model the probability of a subject choosing one of the options (e.g. right) as a function of stimulus strength. We parametrize the psychometric curve as a 4-parameter logistic function:

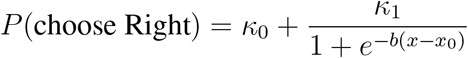

where *x_0_* is the threshold parameter that additively biases the stimulus *x, b* measures sensitivity to the stimulus, *κ_0_* is the left asymptote or left lapse rate and κ_1_ scales the logistic function. Therefore, the right asymptote is given by *κ_0_* + κ_1_ and the right lapse rate itself is given by 1 — (κ_0_+κ_1_). We fit all four of these parameters *{κ_0_, κ_1_,x_0_, b}* to choices generated by either the DDM (Fig 1), rats (Fig 2, 3, 4), or accumulator models adapted to the tasks (Fig 3, 4) using a gradient-descent algorithm (interior-point) to maximize the (Binomial) log likelihood of choices using MATLAB’s constrained optimization function *fmincon. κ_0_* and *κ_1_* were both constrained to lie within the interval [0, 1]. 95% confidence intervals on these parameters were generated using bootstrapping.

### History modulation of psychometric parameters

To summarize the effects of trial history on psychometric parameters we fit independent psychometric curves to choices conditioned on 1-trial back choice-outcome history i.e. following rightward wins (Rw) and leftward wins (Lw). Modulation of the threshold parameter by history was then computed as 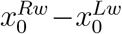. To quantify the modulation of lapse rate parameter by history we first computed the difference in the left and right asymptotes following rightward and leftward wins: 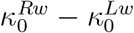 and 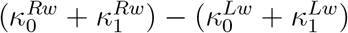 respectively. The net modulation of lapse rates with trial history is given by the sum of these differences: 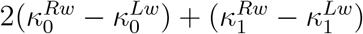.

## Behavioral tasks

### Auditory evidence accumulation task

Rats were trained with previously established protocol (Brunton et al., 2013; Hanks, Kopec, et al., 2015; Erlich, Brunton, et al., 2015; Yartsev et al., 2018) using the BControl system. Briefly, rats were put in an operant chamber with three nose ports. They were trained to begin a trial by poking their nose into the middle port. This initiated two simultaneous streams of randomly-timed discrete auditory clicks for a predetermined duration after a variable delay (0.5-1.3s), one from a speaker to their left and the other to their right. Rats were required to maintain “fixation” throughout the entire stimulus (1.5s), failure to do so led to a violation trial. At the end of the stimulus, rats had to poke towards the side which played the greater number of clicks to obtain a water reward. Stimulus difficulty was varied from trial-to-trial by changing the ratio of the generative Poisson rates of the two click streams. Trial difficulty and rewarded side were independently sampled on each trial.

We analyzed rats which performed greater than 30,000 trials, at 70% or more accuracy. Sessions with less than 300 trials or less than 60% accuracy for either of the choices were excluded. Since rats typically perform this task for many months after having passed the final training stage, to minimize nonstationarities in the data (due to break in training because of holiday closures etc.) and ensure that we are analyzing asymptotic performance, we identified temporally contiguous sessions with stable accuracy by performing change-point detection on smoothed trial hit rate using MATLAB’s *findchangepts* function. The partition with most number of trials was included in the analysis. Since the animals neither made a choice nor received an outcome on violation trials, we ignore them while computing trial-history effects.

### Auditory evidence accumulation task with reaction time reports

To measure rats’ reaction times in addition to choices we modified the auditory evidence accumulation task in two ways. First, we relaxed the “fixation” requirement and instead allowed rats to sample the stimulus for as long as they want. As soon as rats broke fixation by removing their nose from the center port, the stimulus stopped and the rats were required to report their decision by poking into one of the side ports. For any given trial, the time that the rat spent sampling the stimulus was its reaction time. Second, we rewarded rats if they correctly reported the side which had greater underlying Poisson rate rather than the side which played the greater number of clicks. This helped eliminate the trivial strategy of culminating a decision after the first click and having perfect accuracy by simply reporting the side of that click without any need for evidence accumulation.

In practice, we followed the same training protocol as the interrogation task (Brunton et al., 2013) but with the modified reward rule. Once the rats were fully trained on the interrogation protocol we gradually reduced the duration of delay between stimulus onset and trial initiation as well as the fixation period. Most rats maintained high accuracy (>70%) upon this manipulation, if rats performance did not meet this criterion even after a week of training, they were excluded. Rats tended to have worse accuracy early in the session, so we omitted the first 50 trials from our analysis. After the first 50 trials, we confirmed that the accuracy in the first and second halves of the session was comparable.

## Data modeling methods

### Accumulator model

To model subjects choices and RTs, we used accumulation to bound model modified to take into account the discrete nature of evidence in our behavioral tasks (Brunton et al., 2013). In the model, the evolution of accumulated evidence *a(t*) in response to the left (δ_L_) and right (δ_R_) click trains on trial *n* is given by:

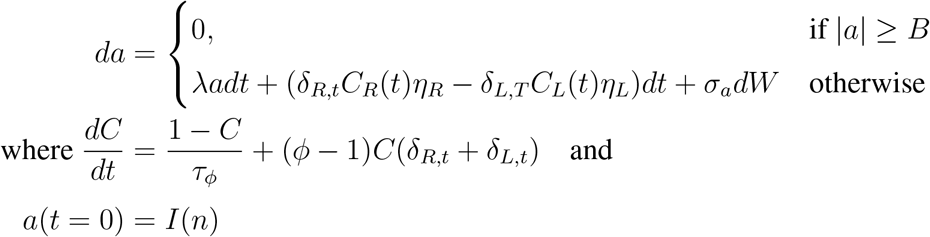

where λ is the inverse time constant of the consistent drift in memory of *a(t). C_R_(t*) and *C_L_(t*) are the magnitudes of each right and left click respectively after undergoing sensory adaptation (with adaptation strength *φ* and adaptation time constant τψ). The sensory noise that accompanies each click is represented by *η_R_, η_L_* which are Gaussian random variables with mean 1 and variance 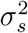. The accumulation variable *a* also undergoes Brownian diffusion through the addition of a Wiener process (*W*) with variance 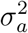. *B* represents the absorbing decision bound that prevents *a(t*) from evolving further, if crossed. The initial value of the accumulator variable *a* varies from trial-to-trial and is set based on exponentially filtered history of previous choices and outcomes (see Methods section on Models of initial state updating). A choice is made by comparing the final value of the accumulator *a*(*T*) to a side bias. A rightward choice is made if *a*(*T*) > bias.

Since the model quantifies noise sources on each trial, it requires estimating the evolution of a noise-induced probability distribution *P(a(t)*). We compute *P(a(t)*) by solving the Fokker-Planck equations that correspond to model dynamics (see Brunton et al. 2013; DePasquale et al. 2021 for numerical methods). The probability of making a rightward choice at the end time-point *T* of a trial, given accumulation model parameters *θ^acc^* is:

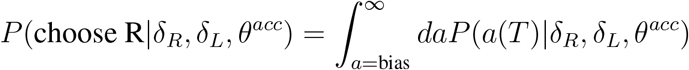

### Models of true lapses

We assume that some fraction of choices *κ* arise from processes extraneous to evidence accumulation such as motor error/exploration or inattention. We parameterize these processes with *θ^lapse^* and refer to them as “true lapses”:

- In the motor error/exploration variant, the probability of making a choice towards the right - when lapsing - is given by *ρ*.

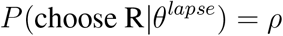
- In the inattention variant (Supp Fig 4C), the subject lapses towards the side favored by the initial state relative to a bias *ρ*. So the probability of a rightward choice due to inattention on trial *n* is:

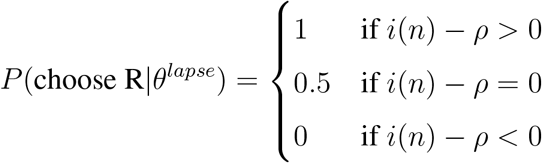
- In the hybrid variant (with motor error and inattention; Supp Fig 6), the probability of lapsing towards right depends on the initial state through a sigmoidal function whose slope *m* (or matching constant) as well as bias *ρ* is a free parameter:

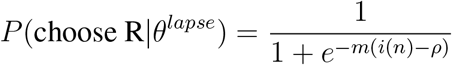

Hence the total probability of making a rightward choice due to accumulation and true lapses is:

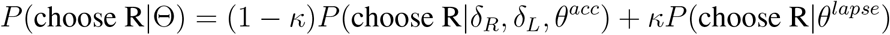

where Θ =*{θ^acc^, θ^lapse^, κ}*.

### Model fitting

The model parameters were fit to individual rats by maximizing the log likelihood of the observed choices of the rat c_obs_, i.e. by maximizing

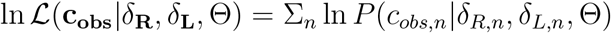

where *n* indexes trials. Constrained optimization was performed in Julia using Optim package. We computed gradients for parameter optimization using a forward-mode automatic differentiation package. The reported maximum likelihood parameters and likelihood values (used for model comparison) are from model fits to the entire dataset. We fit a random subset of 10 rats using 5-fold cross-validation (85% training dataset, 15% test dataset) but this yielded very similar maximum likelihood parameters and virtually identical test and training log-likelihoods. Hence, to save on computing time we fit the different model variants to each rat’s entire dataset. This agreement between test and training likelihoods is likely due to the large number of trials in our dataset and the modest number of parameters in our model.

### Simultaneous modeling of choices and RTs

In decision-making tasks, observed reaction times (RTs) are often thought of as comprised of stimulus sampling or decision times (DTs, the time it takes for the subject’s accumulated evidence to hit the bound) and non-decision related processing times (NDTs). In our datasets we observed that reaction times tended to be slower following incorrect trials and that they grew longer over the course of a session. These effects could be isolated just to RTs and were not observed in choice behavior. To model these trends we conceptualize non-decision times as arising from a separate drift diffusion process whose drift *ν* is additionally modulated by current trial number n and previous trial’s outcome. These non-decision time drift-diffusion processes terminate when the bound *ω* is hit. We assume that the non-decision times for each choice *k* ∈ {*L, R*} have independent bounds (*ω_k_*) and drifts (*ν_k_*). So the non-decision times for a trial n are samples from the following Wald or Inverse Gaussian (*IG*) distribution:

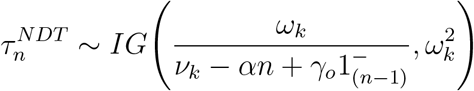

where *k* ∈ {*L, R*} and 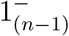 is an indicator function which is 1 if the previous trial was incorrect and is 0 otherwise. *α* parameterizes the impact of trial number on NDTs and *γ_o_* parameterizes the impact of previous trial’s outcome on current trial’s NDT.

We fit the model by maximizing the joint log likelihood of the observed choices and RTs. For any given trial, we can compute the likelihood of observing a particular reaction time *RT_obs_* and choice *c_obs_* due to accumulation by marginalizing over possible decision or bound hitting times *τ_c_obs__* for the observed choice:

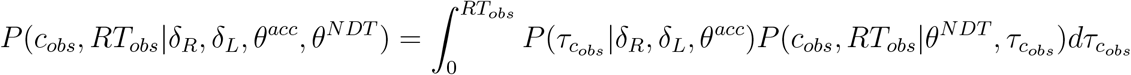

On true lapse trials, RTs were assumed to arise from NDTs alone and therefore the joint likelihood due to accumulation and true lapses is given by:

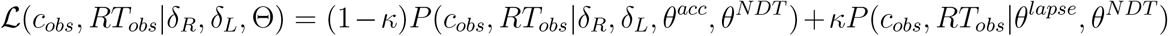

where Θ = {*θ^acc^, θ^N DT^, θ^lapse^, κ*}.

## Acknowledgements

We thank members of the Brody lab for experimental support and helpful feedback throughout the project especially Adrian Bondy, Thomas Luo, Emily Dennis, Tyler Boyd-Meredith, and Ahmed El-Hady. We also thank Jovanna Teran and Brody lab technicians for assistance with rat training. We are grateful to Sashank Pisupati, Jonathan Cohen, Sebastian Musslick, Jonathan Pillow, and Ilana Witten for helpful discussions at various points during the project. This work was supported by NIH grant R01MH108358 to CDB.

## Supplementary materials

**Supplementary Figure 1:**
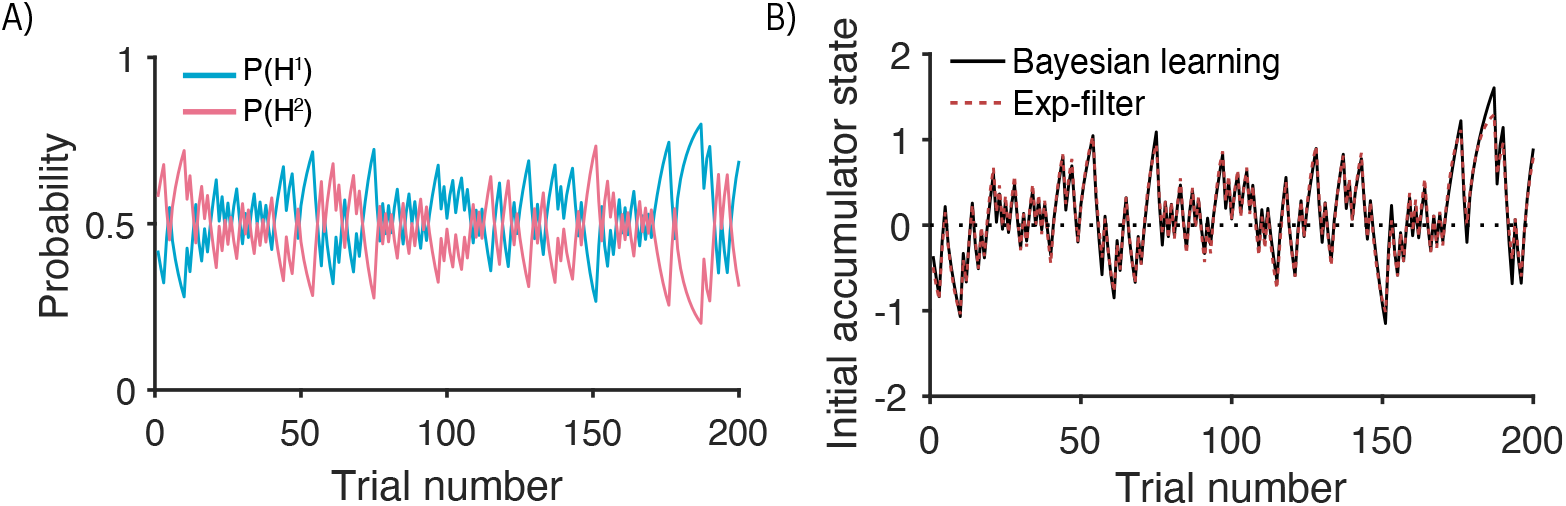
Exponential filtering for initial state setting approximates Bayesian prior updates under assumptions of non-stationarity. Example of a mis-belief in a non-stationary prior. Traces represent belief about prior probability of two hypotheses *H*^1^ and *H*^2^ inferred from a random sequence of trials drawn from a stationary symmetric prior, under the misbelief that the prior is occasionally undergoing unsignalled jumps. Such an assumed generative model is often referred to as the Dynamic Belief Model (DBM; Yu and Cohen 2009). **B**: Initial state updates corresponding exactly to the fluctuating prior beliefs in (A) that emerge from Bayesian learning (black line), plotted against approximate initial states derived from exponential filtering (dotted red line) of past choices and outcomes. The exponential filter provides a good approximation of exact Bayesian updates, while being more expressive and flexible to capture the possibility of other generative models and corresponding update rules.

**Supplementary Figure 2:**
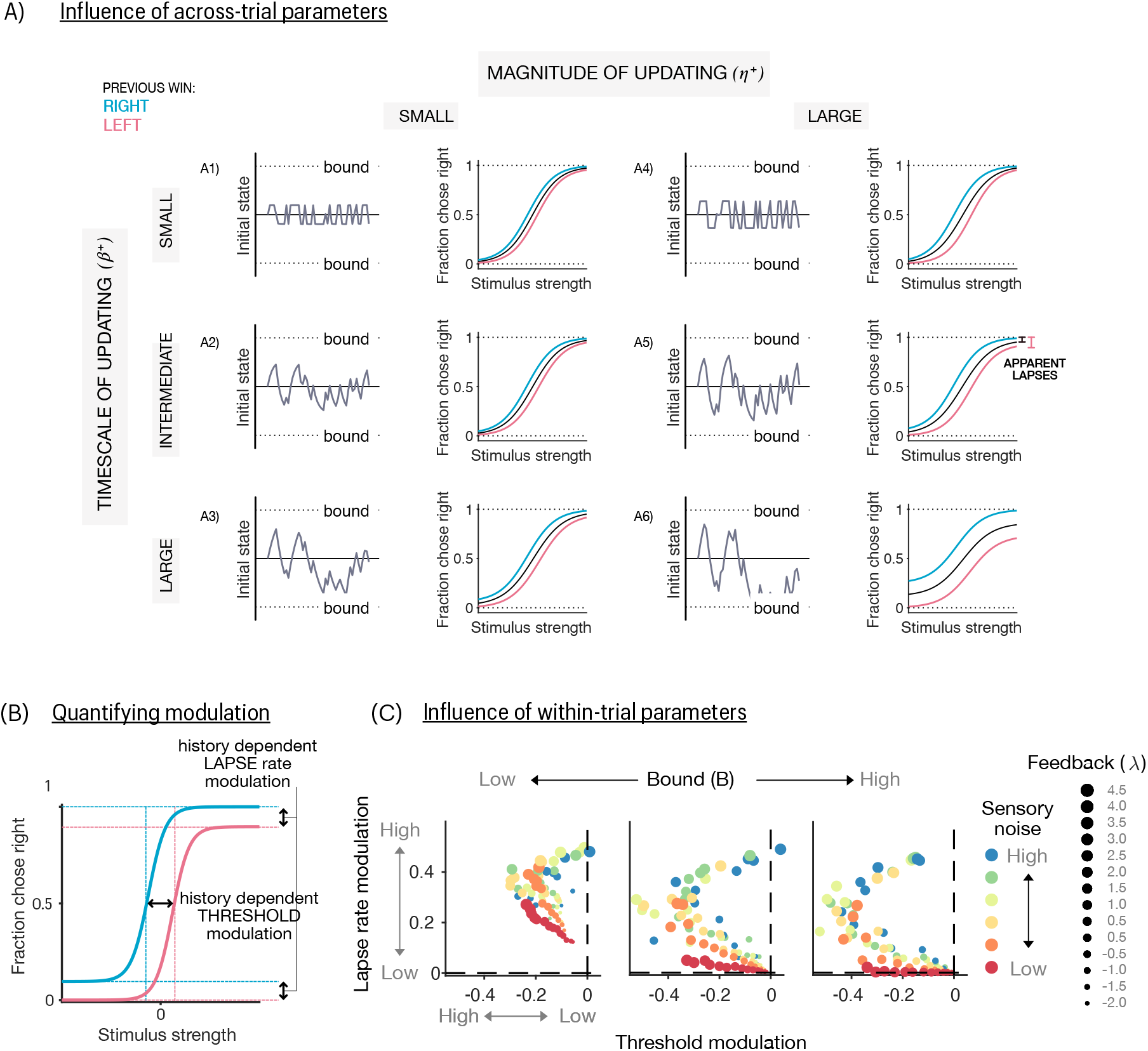
Influence of within- and across-trial parameters on history-dependent modulation of biases and lapses in the psychometric function. **(A)** Influence of across-trial parameters on history-dependent modulation: Effects of varying magnitude of trial-by-trial updating (η - columns) and timescale of updating (*β* - rows) on initial state trajectories (gray lines) and psychometric curves (black - conditioned on all previous wins, blue - previous right wins, pink-previous left wins). (Top row) Small timescales of updating lead to fast fluctuations in initial states, and mostly horizontal shifts in psychometric curves with trial history, for both small and large magnitudes of updating (A1 and A4). (Middle row) Intermediate timescales of updating lead to slower fluctuations in initial state that have a cumulative effect across trials. For large magnitudes of updating (A5) these can give rise to apparent lapses (black intervals) as well as history-dependent modulation of these lapses (pink intervals). (Bottom row) Long timescales of updating lead to stronger cumulative initial state biases across trials, yielding apparent lapses and lapse rate modulations even for small magnitudes (A3). When combined with large magnitudes of updating (A6) lead to initial states that sometimes exceed the bounds, leading to a combination of apparent lapses (initial states within bounds) and deterministic, stimulus-independent decisions (initial states outside bounds). **(B)** Quantifying modulation of psychometric properties: The difference between psychometric curves conditioned on previous wins (blue) or losses (pink) can be quantified using two metrics - the horizontal distance between the midpoints of psychometric curves (“threshold modulation”) and the vertical distance between its asymptotes (“lapse rate modulation”) **(C)** Effects of varying the parameters of the within-trial drift diffusion model (DDM) on history-dependent threshold (x-axis) and lapse rate modulations (y-axis). Colors denote levels of sensory noise, size of dots denote values of the feedback parameter of the DDM. The feedback parameter determines if the accumulation is leaky (λ < 0, ignores early evidence), perfect (λ= 0, uses all evidence) or impulsive (λ> 0, ignores later evidence). Plots from left to right are for low, intermediate and high values of bound respectively. High bounds predominantly give rise to threshold modulations, however high positive values of feedback and higher levels of sensory noise additionally produce lapse rate modulations. Lapse rate modulations are dramatically increased by lower bounds for many different values of feedback and by higher values of sensory noise.

**Supplementary Figure 3:**
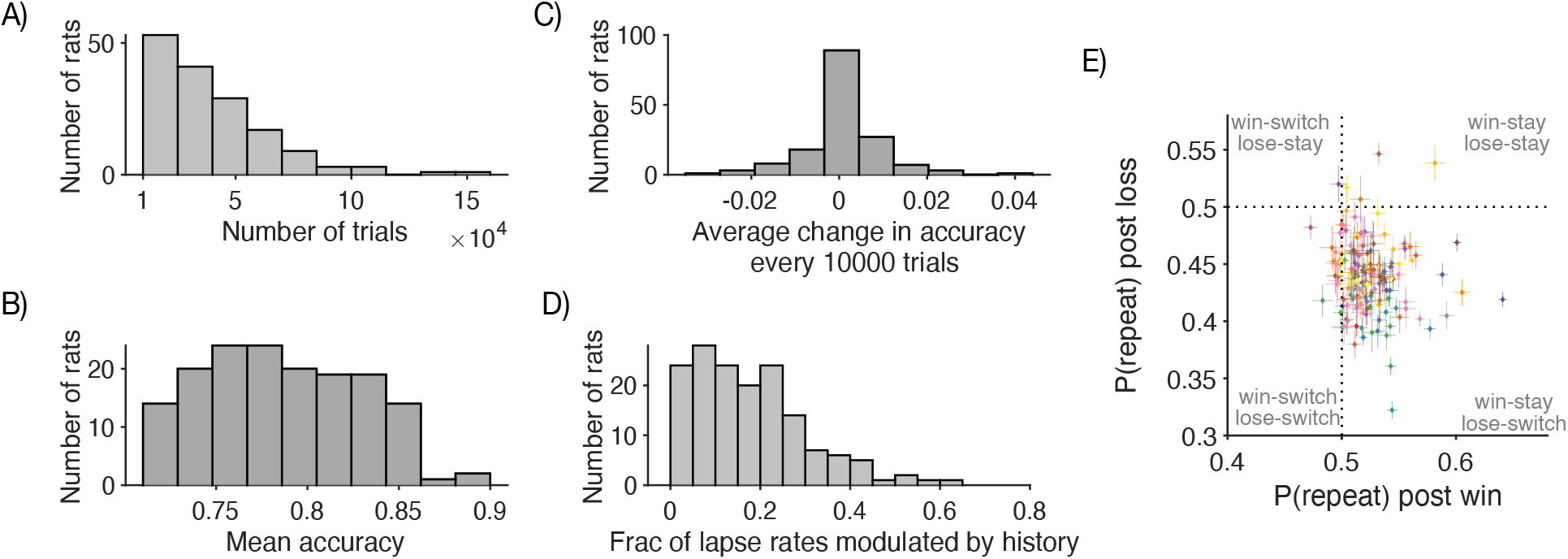
Performance measures across the rat dataset. **(A)** Histogram of trial counts for all rats in the population. Most rats completed on the order of 1e^4^ trials. **(B)** Histogram of mean accuracy showing that rats showed good performance on the task (mean accuracy ±SD: 0.79 ±0.04). **(C)** Average change in mean accuracy every 10000 trials, showing that rats’ performance was stable over time, reflecting asymptotic behavior rather than task acquisition. **(D)** Histogram of history-modulated lapse rates as a fraction of total lapse rates. A sizeable portion of the population had non-zero fractions, suggesting that history-dependence could potentially account for substantial lapse rate variance. **(E)** Scatter comparing repetition bias following wins and losses. Each point is a rat, error bars are Wilson binomial CIs. Most of the population occupied the bottom right quadrant, showing a “win-stay, lose-switch” bias i.e. repetitions following wins and alternations following losses.

**Supplementary Figure 4:**
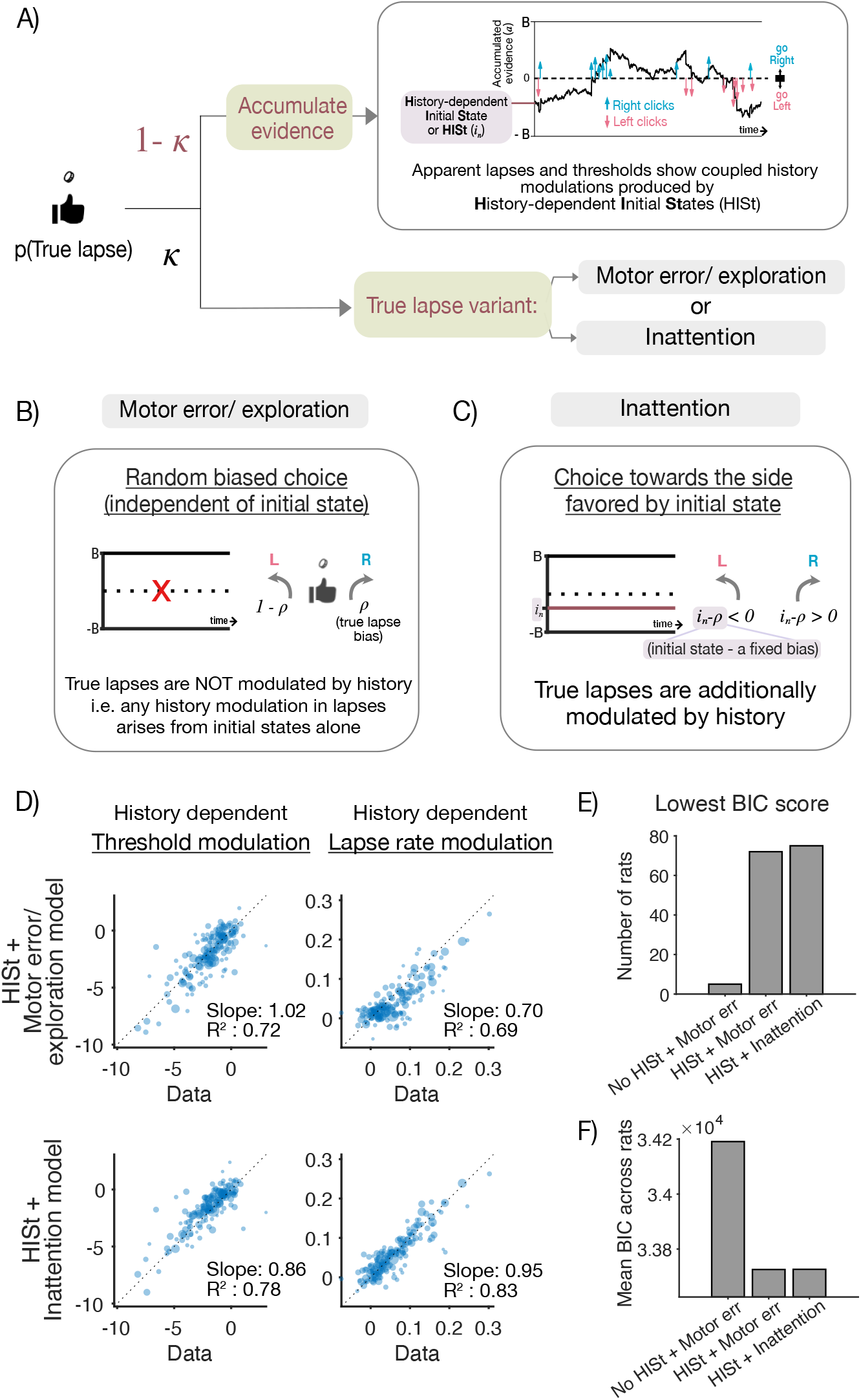
Two variants of the accumulator with HISt model with different kinds of true lapses perform equally well. **(A)** Schematic of accumulator with HISt (top), which produces apparent lapses and thresholds that are both modulated by history, with two variants of true lapses (bottom) - those due to motor errors/exploration, and those due to inattention. **(B)** Motor error/exploration variant, that occasionally chooses a random action with some bias (ρ) irrespective of the initial state, reflecting an error in motor execution or random exploration. This model produces true lapses that are not modulated by history, such that any history modulations arise from HISt alone. **(C)** Inattentional variant, that occasionally fails to attend to the stimulus, and relies on the initial state to make an informed, deterministic decision based on the difference between the initial state and a bias (ρ). In this model, true lapses are also modulated by history in addition to apparent lapses and thresholds. **(D)** Individual differences in history effects captured by different models: History modulations of threshold (left) and lapse rate (right) parameters measured from psychometric fits to the raw data (x-axis) v.s. model predictions (y-axis). (Top): Motor error/exploration model has no history dependence in true lapses, yet captures a majority of the variance in both threshold and lapse rate modulations [R^2^ = 0.72 (threshold parameter), *R*^2^ = 0.69 (lapse rate parameter)], and shows good correspondence with both parameters, as evidenced by the points lying close to the unity line [slope= 1.02 (threshold parameter), slope = 0.70 (lapse rate parameter)]. This suggests that these modulations can be captured by the comodulations in apparent lapses and thresholds produced by HISt. (Bottom): same as Top but for Inattention model. The inattention model allows true lapses to additionally depend on history, and captures slightly more variance in history modulations [R^2^ = 0.78 (threshold parameter), R^2^ = 0.83 (lapse rate parameter)]. However, it does so at the expense of correspondence with thresholds [slope= 0.86 (threshold parameter), slope = 0.95 (lapse rate parameter)]. This marginal improvement suggests that comodulations in thresholds and lapse rates largely reflect apparent lapses arising from HISt, rather than true lapses of either kind. **(E)** Distribution of best fitting model variants for individual rats: Overall bar height for each variant denotes the total number of rats for which that variant scored the lowest BIC score. Inattention variant won in marginally more rats than motor error (inattention: 75/152 rats, motor error: 72/152 rats). **(F)** Population model comparison using mean BIC score across rats. Lower scores indicate better fits. Scores are comparable across variants, marginally favoring motor-error over inattention (Mean BIC score for motor error/exploration: 33725.64, inattention: 33726.25).

**Supplementary Figure 5:**
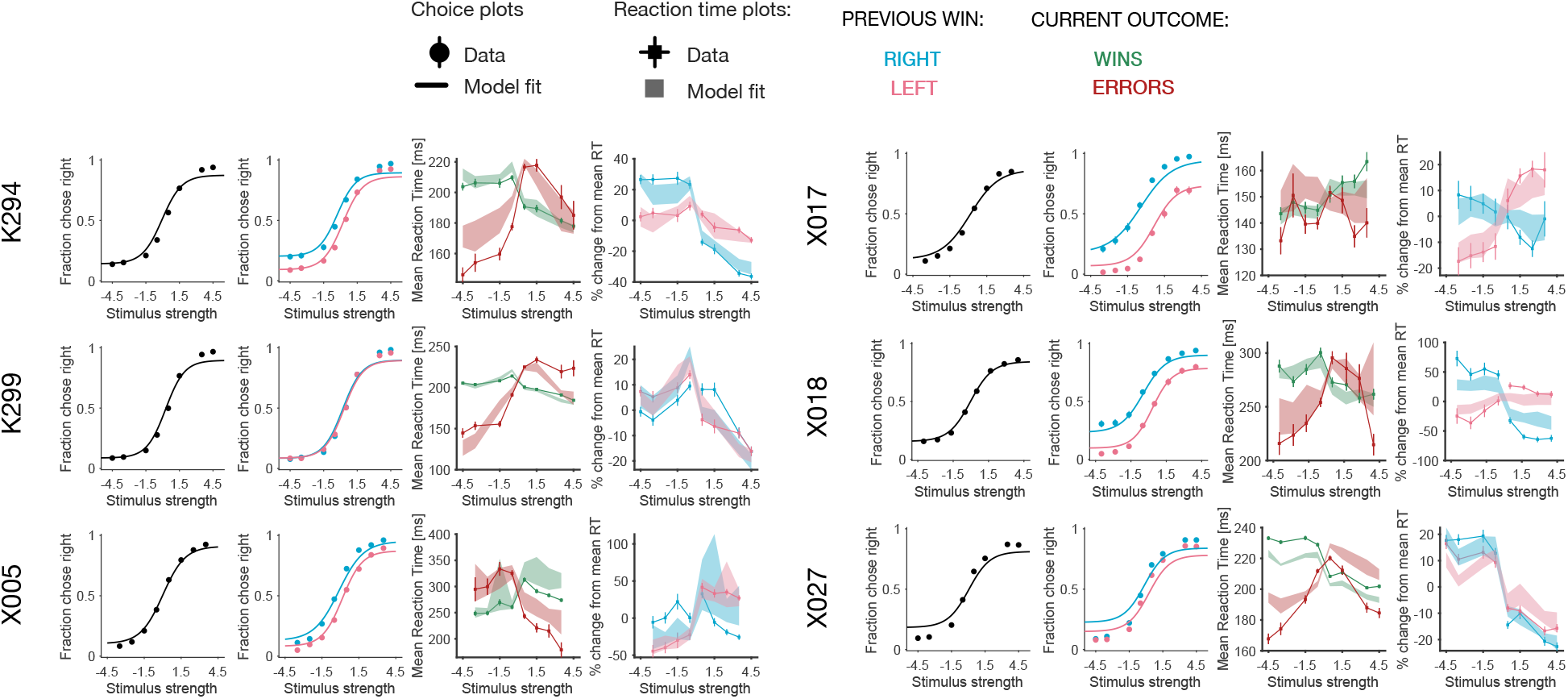
Fits of the accumulator model with history-modulated initial states (and additional true lapses arising from motor error) to choices and reaction times of individual rats. Each horizontal set of 4 panels shows fits to an individual rat, and each of the 4 columns depicts a different behavioral measure summarizing choices (first column, psychometric curve), reaction times (third column, win RTs in green and error RTs in red), and history modulations in choices/reaction time (second/fourth column, psychometric curves/RTs conditioned on previous right wins (blue) or left wins (pink)). Data represented by points (circles: choices, squares: reaction times) and model fits represented by lines (choices) or shaded bars (reaction times, thickness represents 95% bootstrap prediction intervals).

**Supplementary Figure 6:**
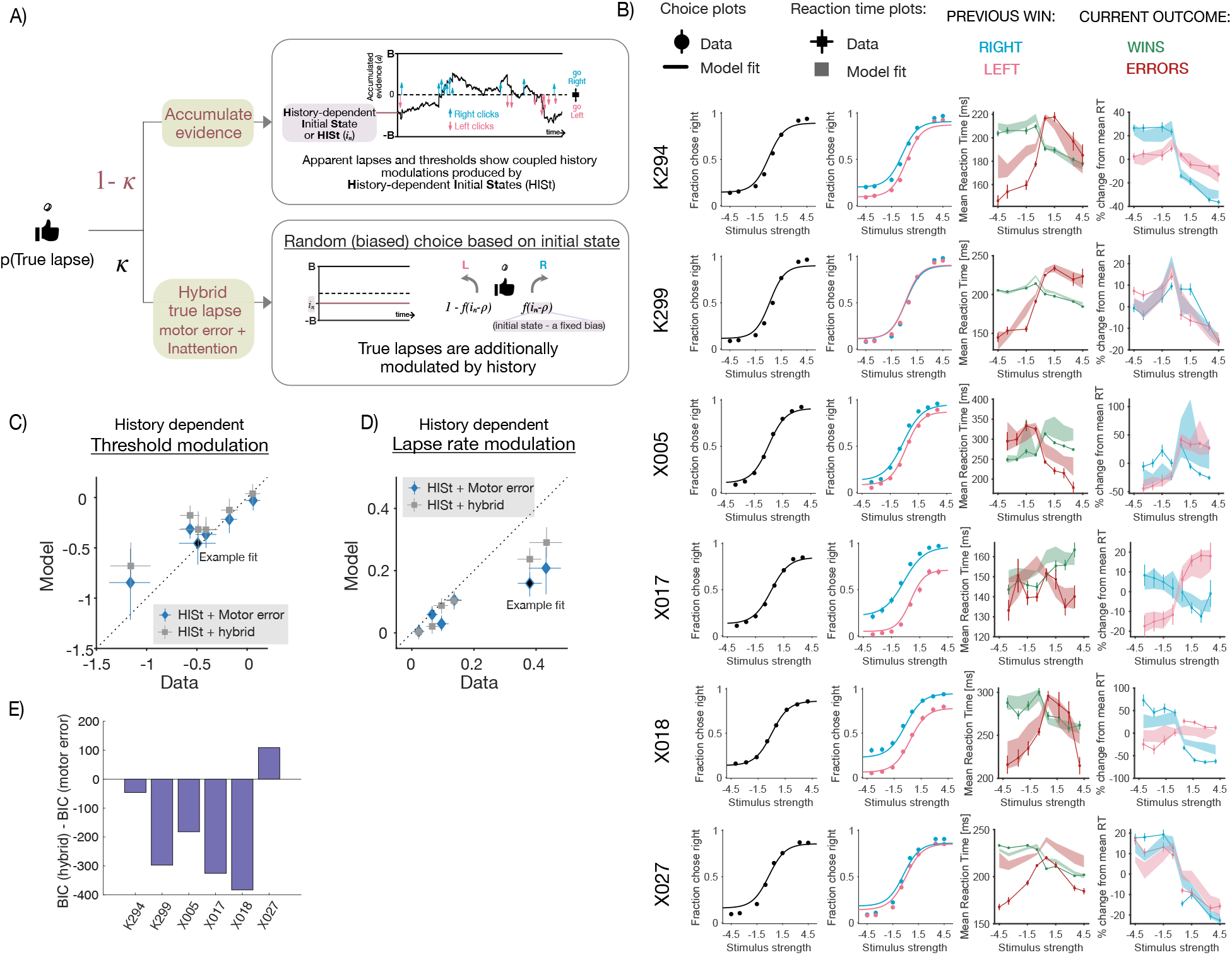
Accumulator model with HISt and true lapse variants fit to the RT dataset. **(A)** Flexible true lapse variant of the accumulator model with HISt, capable of producing both motor errors and inattentional true lapses: in this model, the subject chooses stochastically on true lapse trials, with a probability given by a sigmoidal function of the initial state (such a strategy is often called probability matching and although suboptimal, has found empirical support in many perceptual tasks e.g. Mamassian et al. 2002). The slope parameter of the sigmoid which is fit to the data – when this slope parameter goes to infinity, this model picks deterministically based on the initial state, similar to the previous inattention model. On the other hand when the slope goes to zero, choices on true lapse trials are no longer dependent on initial states, reducing to the motor error/exploration model, with intermediate parameters interpolating between these two extremes. This “hybrid true lapse” variant of the model can flexibly include many kinds of true lapses. **(B)** Fits of the hybrid model to individual rats in the RT dataset, showing choice, RT and history measures similar to Supp Fig. 5.**C-D** History modulations in psychometric thresholds (C) and lapse rates (D) for motor error (blue) and hybrid (grey) models. Once again, allowing for the possibility of history-modulated true lapses slightly improves correspondence to lapse rate modulations, at the cost of threshold modulations. **(E)** Difference in BIC scores between the hybrid and motor error models, for individual rats in the reaction time dataset. Negative values indicate that the more flexible hybrid model won, as is the case for most rats.

## Footnotes

1 In tasks where the reliability of incoming evidence (controlled by stimulus strength) varies from one trial to the next, it has been shown that ideal observers should have time-varying bounds on the posterior (Drugowitsch, Moreno-Bote, et al. 2012. However under certain circumstances, stationary bounds over the summed stimulus have been shown to implement close-to-optimal collapsing bounds on the posterior, which is the regime we assume here for simplicity.

2 For a treatment of non-stationary likelihood functions which yield variability in drift rate, see (Drugowitsch, Mendonça, et al., 2019; Mendonça et al., 2020)

3 Note that this is the case if the agent assumes that a shift in the prior over stimulus categories maps onto an overall shift in the prior over stimulus difficulties — see (Drugowitsch, Mendonça, et al., 2019) for a detailed treatment

